# REMY: A platform for the rapid interrogation of epigenome modifications on yeast

**DOI:** 10.1101/2021.02.24.432679

**Authors:** Alison C. Waldman, Balaji M. Rao, Albert J. Keung

## Abstract

Histone proteins are decorated with a combinatorially and numerically diverse set of biochemical modifications. Here we describe a versatile and scalable platform termed **R**apid interrogation of **E**pigenome **M**odifications using **Y**east surface display (REMY), which enables efficient characterization of histone modifications without the need for recombinant protein production. As proof-of-concept, we first used REMY to rapidly profile the histone H3 and H4 residue writing specificities of the human histone acetyltransferase, p300. Subsequently, we used REMY to screen a large panel of commercially available anti-acetylation antibodies for their specificities, identifying many suitable and unsuitable reagents. Further, use of REMY enabled efficient mapping of the large binary crosstalk space between acetylated residues on histones H3 and H4, and uncovered previously unreported residue interdependencies affecting p300 activity. Our results show that REMY is a useful tool that can advance our understanding of chromatin biology by enabling efficient interrogation of the complexity of epigenome modifications.

## Introduction

Chromatin is numerically and combinatorially complex with over 60 distinct biochemical histone modifications known to date. This complexity presents considerable challenges to understand the regulatory logic and function of chromatin as well as to simply map the properties of individual chromatin components. To address this complexity, new technologies have recently been developed, including libraries of DNA-barcoded nucleosomes that can rapidly map the binding specificities of protein domains to diverse histone modification patterns in vitro^1^, genetic approaches such as a yeast strain library that was used to assess the impact of histone H3 and H4 mutants on cell physiology^2^, and large histone peptide arrays used to map the binding partners of different bromodomain families^3^.

Despite these and other advances, there remain several important gaps in our capabilities. In vitro methods provide exquisite control over residue specificities yet require recombinant production that often is refractory for many proteins or chemical syntheses and ligations that can become laborious to scale^4^. These limitations are particularly acute in investigating enzymatic activities, with no current high throughput and easy to implement approach that can assess the specificities of epigenome ‘writers’ for their histone substrates.

Here we describe a platform for the efficient interrogation of epigenome modifications using yeast surface display (REMY). This platform bypasses the need for recombinant protein production or chemical ligation, preserves the ability to assess residue specificities as with in vitro biochemical assays, and is able to probe enzymatic activities of histone writers. In REMY, the histone tail is co-expressed as a C-terminal fusion to the yeast cell wall protein Aga2p in the conventional yeast surface display system^5^, along with the epigenome writer. Both the histone tail and the epigenome writer are targeted to the endoplasmic reticulum, where the histone tail is modified by the writer, and results in cell surface display of the modified histone tail (**Fig. 1A**). Using REMY, we demonstrate histone acetylation by the histone acetyltransferase (HAT) p300 and map the specific histone H3 and H4 residues that are modified by p300. We further show how REMY provides a rapid way to rigorously assess the quality of commercial histone antibodies. Finally, by comprehensively mapping all binary crosstalk interactions between residues on histones H3 and H4, we demonstrate the scalability of this system that is dependent only on the construction of DNA expression libraries, not on recombinant protein production.

**Figure 1.**
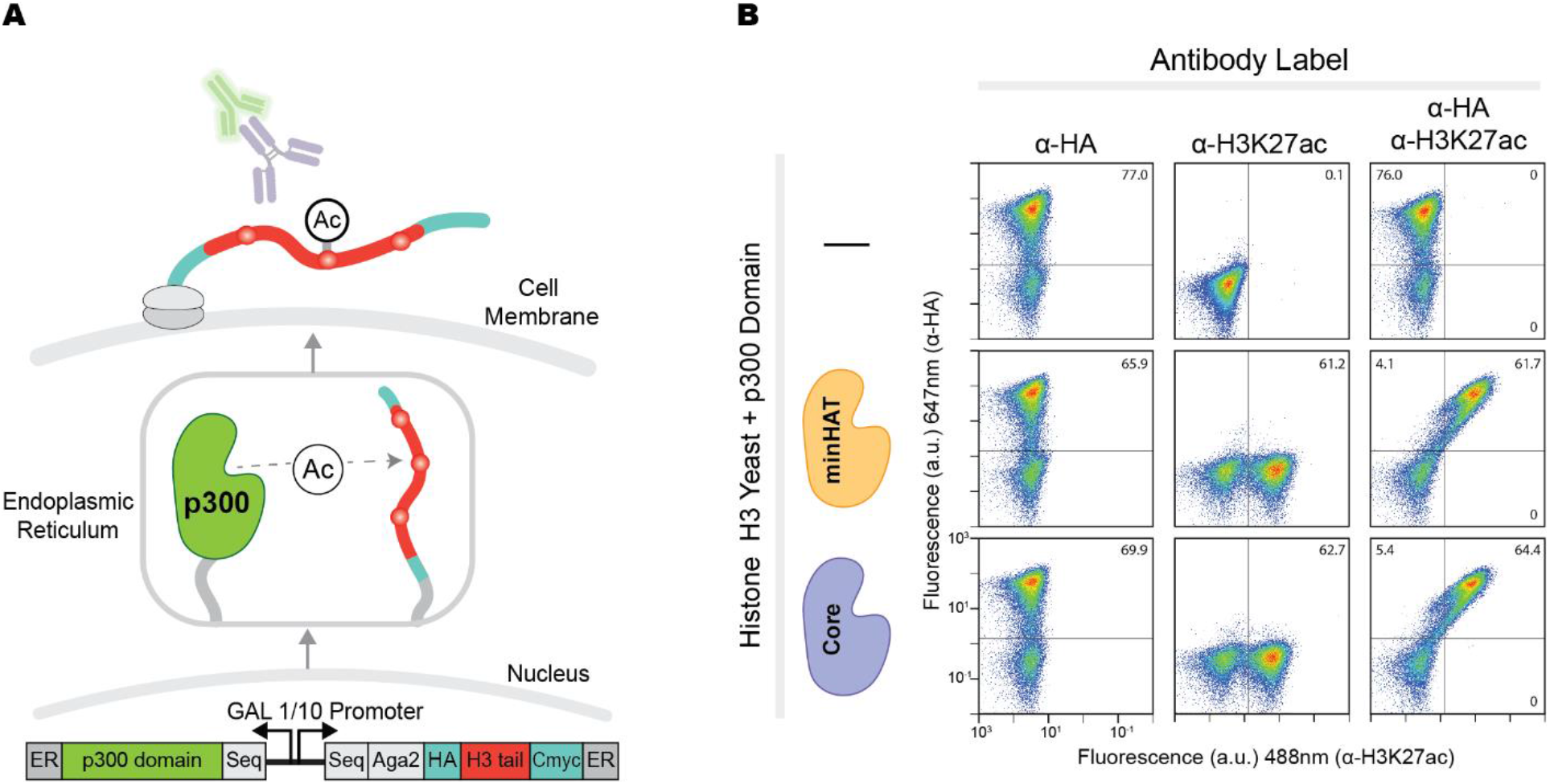
REMY efficiently acetylates and displays histone peptides. (A) A schematic of the endoplasmic sequestration and Aga1p-Aga2p yeast surface display method. The histone peptides have N-terminal HA tags and C-terminal Myc tags that allow for quantification of their expression and display on a single cell basis. (B) Flow cytometry confirmation that the histone tails are displayed and only acetylated when a p300 writer domain is present. The HA tag measures the expression level of the histone H3 tail. The double labeled plots show the high correlation between histone display levels and acetylation when p300 is present. Fluorescent gates were created based on unlabeled cells of the correspond yeast strain per each sample (not shown). The percentage of cells above each gate is recorded in top corner of each plot. All plots contain 80,000 cells.

## Results

### REMY efficiently acetylates and displays histone peptides

REMY uses yeast endoplasmic reticulum (ER) sequestration screening (YESS)^6^ to colocalize an epigenome enzyme with a histone tail peptide, allowing the peptide to be post-translationally modified. The histone peptide fused to the Aga2p protein, and the epigenome writer are transcribed simultaneously by an inducible, bidirectional GAL1/10 promoter (**Figure 1A**). By tagging with ER sequestration and retention sequences, both proteins enter the ER and temporarily dock into the interior ER membrane. Here, the writer modifies the histone due to their high effective local concentration and proximity. Both proteins then continue through the secretory pathway where the writer is secreted into the supernatant and the modified histone is tethered to the yeast surface through the disulfide bond between the Aga1p and Aga2p proteins.

As a proof of principle, we chose the human histone acetyltransferase (HAT) E1A binding protein p300 (p300 or KAT3A) as the writer and histones H3 and H4 as the substrates. p300 is a well-characterized HAT that acetylates histones and plays a significant role in chromatin regulation *in vivo*^7–9^. Acetylation of lysine residues on histones H3 and H4 has been closely tied to activation of gene transcription^10–13^. Further, transcription strength may be regulated by the number of acetylated residues on a histone tail^14^ or preferential acetylation of specific lysine residues^13,14^.

We first investigated the acetylation of H3K27 in REMY; H3K27 has been widely reported to be acetylated by p300^15,16^. We used two different domains of p300, the core catalytic domain (Core, aa1048-1664)^17^ and the minimal HAT domain (minHAT, aa1284-1669)^18^. Labeling with antibodies against H3K27ac and HA showed that the acetylation signal (H3K27ac) correlated strongly with the amount of histone peptide displayed (HA) (**Figure 1B**). No acetylation was detected when p300 was absent. These results confirmed that histone peptides could be modified and displayed as cell surface fusions using REMY.

### REMY can assess specificity of anti-acetylation antibodies

Validation of the advertised specificities of commercially available antibodies is a significant challenge for the chromatin biology field^19^. We investigated if REMY could provide a general platform to test the specificity of antibodies against histone modifications. To independently assess the antibody specificities of histone H3 and H4 acetylation antibodies for our specific study, we first used chemically defined histone peptides immobilized on the surface of yeast using the streptavidin-biotin linkage. Briefly, a modified streptavidin monomer (mSA^20–22^) was expressed as a cell surface fusion using yeast surface display. We purchased biotinylated (but otherwise unmodified) histone H3 and H4 peptides as negative controls and biotinylated, singly-acetylated peptides as positive controls (**Supplemental Table 1**). These peptides were linked to the surface of the mSA-displaying yeast. The specificities of an assortment of anti-acetylation antibodies targeting each of the available lysine residues on the N-terminal tails of histones H3 and H4 (**Supplemental Table 2**) were tested by the ability of the antibody to specifically detect the appropriate acetylated peptides by flow cytometry (**Figure 2**). In parallel, we investigated whether REMY could be used to assess the specificity of these antibodies. Towards this end, yeast cells expressing the histone peptides in the presence or absence of the p300 domains were labeled with the selected antibodies. Yeast cells expressing mutant histones with single arginine mutations at each of the lysine residues on the histone H3 and H4 tails were also generated (**Supplemental Table 3**), and served as negative controls for the antibodies. The arginine mutant controls provided confidence that the antibodies were not binding other residues non-specifically that might be acetylated by p300 (**Figures 2B & 3**). Collectively, we confirmed the specificity of a set of primary antibodies against H3K4ac, H3K9ac, H3K18ac, H3K27ac, H3K36ac, H4K5ac, H4K8ac, H4K12ac, H4K16ac, and H4K20ac (**Figure 2A**), with both approached yielding similar results (**Supplemental Table 2**). However, despite testing multiple commercially available antibodies, we were unable to identify appropriate antibodies for H3K14ac and H3K23ac (**Figure 2B**). These results show that REMY is a useful tool to efficiently assess the specificity of antibodies putatively recognizing histone modifications at specific residues. Antibodies found to be specific using both approaches were used in further experiments in this study.

**Figure 2:**
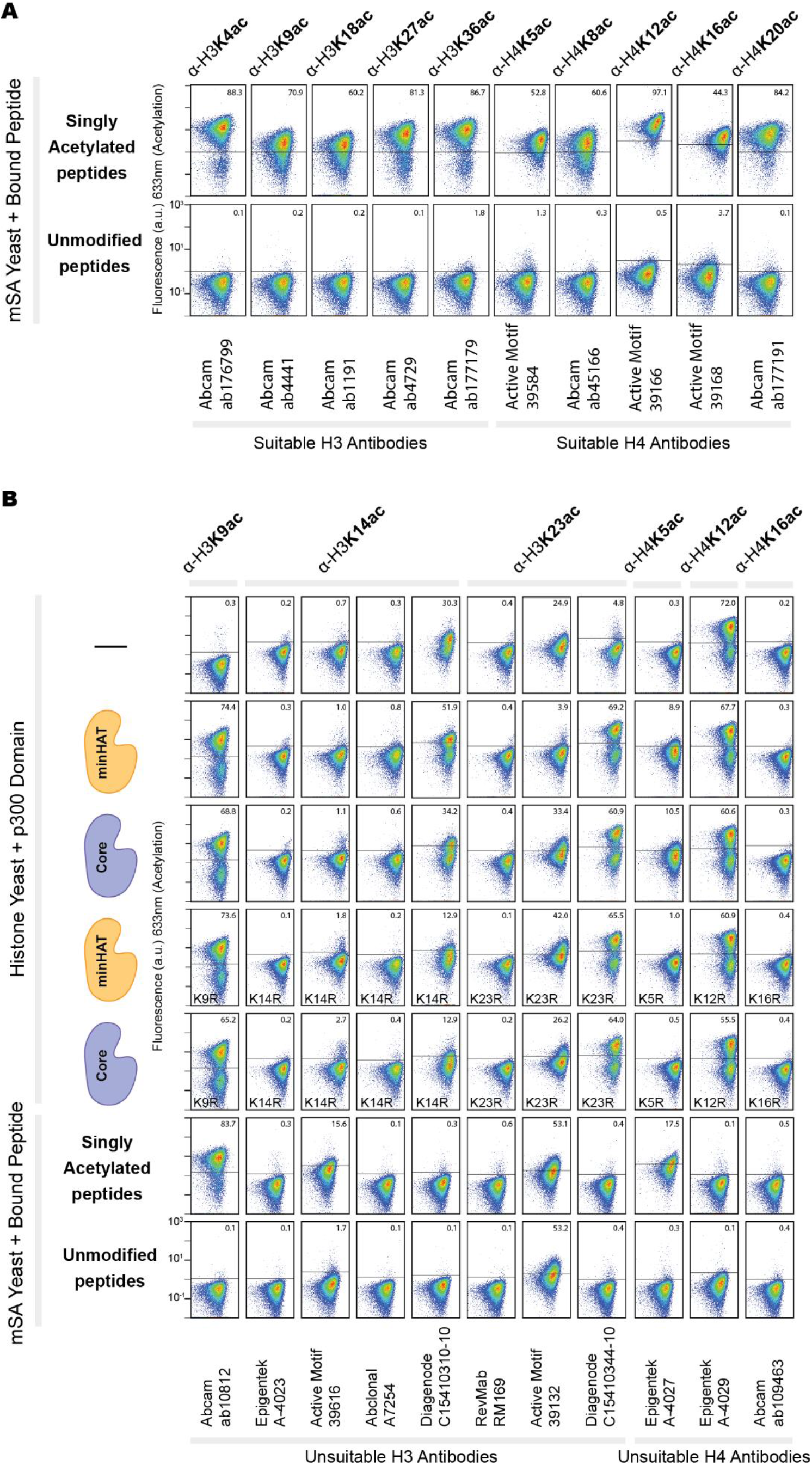
REMY can assess specificity of anti-acetylation antibodies. (A) Primary antibodies against H3K4ac, H3K9ac, H3K18ac, H3K27ac, H3K36ac, H4K5ac, H4K8ac, H4K16ac, and H4K20ac that were selected to map the residue specificities of p300 domains (Supplemental Table 2, bolded). These antibodies showed strong binding to their corresponding singly acetylated peptide when bound to the strep-displaying yeast and showed no binding to yeast displaying an unmodified histone peptide. (B) Anti-acetylation antibodies not suitable for residue mapping. We were not able to identify suitable antibodies against H3K14ac nor H3K23ac. 2E6 yeast cells were labeled with 1 μM peptide for each sample. Fluorescent gates were created based on unlabeled cells of the corresponding yeast strain per each antibody (not shown). The percentage of cells above each gate is recorded in top corner of each plot. The streptavidin monomer has an N-terminal HA tag and a C-terminal FLAG tag which were labeled to confirm high expression levels of mSA (data not shown). All plots contain 80,000 cells.

**Figure 3.**
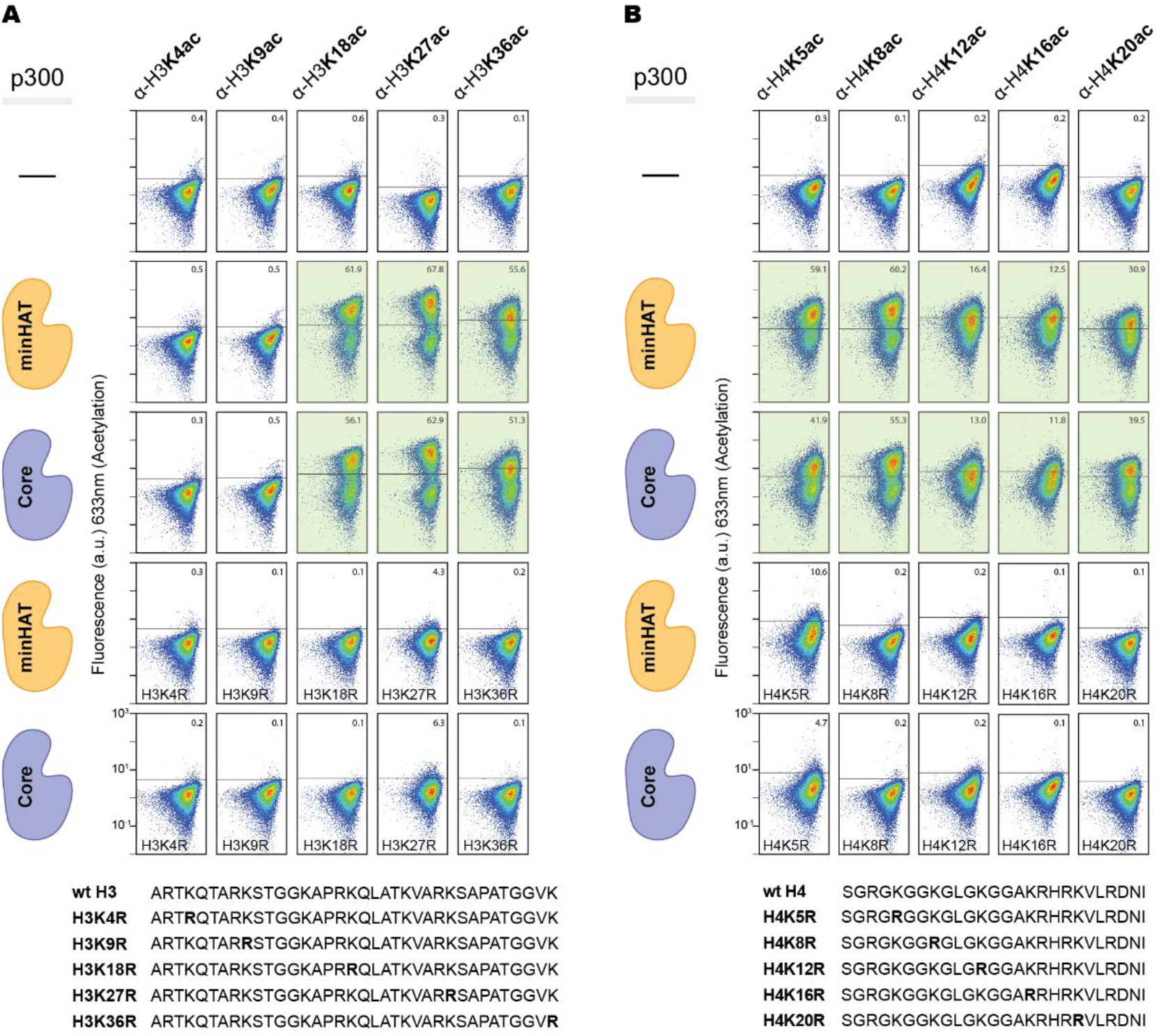
REMY uncovers the residue specificities of p300. The y-axes display the fluorescence due to binding of the corresponding anti-acetylation antibodies for the N-terminal tails of (A) histone H3 and (B) histone H4 expressed with or without p300 domains. The plots highlighted in green indicate which residues were acetylated by p300. The p300 domain expressed in each sample is indicated on the left. Lack of antibody binding on the different arginine-mutated histones indicate that the antibodies are not binding to acetyl groups deposited on off-target lysines. Fluorescent gates were created based on unlabeled cells of the corresponding yeast strain per each antibody (not shown). The percentage of cells above each gate is recorded in top corner of each plot. All plots contain 80,000 cells and these acetylation trends were observed for three biological replicates of each sample.

### REMY uncovers the residue specificities of p300

Mapping the residue specificities of epigenome writers is an important step in ultimately understanding their regulatory roles and functions. Using the set of suitable antibodies we identified, we mapped the residue specificities of the minimal HAT and Core domains of p300 for the N-terminal tails of both histones H3 and H4 using REMY and flow cytometry. We determined that the p300 minimal HAT and Core domains both acetylated histone H3 at K18, K27, and K36 and acetylated histone H4 at K5, K8, K12, K16 and K20 (**Figure 3**). We did not observe acetylation at H3K4 or H3K9. Notably, these acetylation patterns match what has been previously observed using biochemical assays^17,23–28^ (**Table 1**). Furthermore, mutating each specific residue to arginine ablated labeling by the matching antibody. These results show that REMY can be used to assess residue specificities of epigenome writers.

**Table 1:** Prior studies that reported histone residues acetylated by p300.^17,23–28^

| Publication | H2A and H2B residues | H3 residues | H4 residues |
| --- | --- | --- | --- |
| Schiltz et al 1998 | p300 acetylates all sites on H2A and H2B known to be acetylated in bulk chromatin - H2AK5 and H2BK5, K12, K15, K20 | p300 prefers H3K14ac, H3K18ac over H3K4ac and H3K23ac. p300 did not acetylate H3K9 or H3K27 | prefers H4K5ac, H4K8ac. Saw acetylation at H4K5, K8, and low levels of acetylation at H4K12, K16 |
| McManus et al 2003 | p300 acetylates H2BK5, K12, K15, K20 in COS-7, HeLa, and IM cells | p300 acetylates H3K14, not H3K9 in COS-7 cells. p300 acetylates H3K9 and H3K14 in HeLa and IM cells. | p300 acetylates H4K5 in H4K8 in COS-7, HeLa, and IM cells |
| Kouzarides 2007 | CBP/p300 acetylates H2AK5, H2BK12, and H2BK15 | CBP/p300 acetylates H3K14 and H3K18 | CBP/p300 acetylates H4K5 and H4K8 |
| Bedford et al 2010 |  | CBP/p300 is responsible for global H3K18ac and H3K27ac in mouse fibroblast cells |  |
| Henry et al 2013 |  | p300 acetylates H3K9, K14, K18, K23. very low levels of H3K27ac | p300 acetylates H4K5, K8, K12, and K16 |
| Dancy and Cole 2015 |  |  | p300 prefers H4K5ac and H4K8ac. p300 knockout cells showed reduced acetylation at H4K5, K8, K12, and K16. |
| Hilton et al 2015 |  | p300 Core domain acetylates H3K27 |  |

### REMY reveals crosstalk between modified histone residues

We used REMY to map crosstalk between modified histone residues, specifically the effect of histone modification at each specific residue on the enzymatic activity of p300 for all other residues. This form of crosstalk has been observed in several systems but is difficult to map comprehensively. For example, acetylation of the fourteenth lysine on histone H3 (H3K14ac) has been shown to enable the Rtt109-Vsp75 complex to acetylate the 56^th^ lysine on the same histone (H3K56ac)^29^; when histone H3 is not pre-acetylated at the fourteenth residue, H3K56 is not acetylated. Additionally, some histone modifications have been found in pairs on histone tails, such as H4K5/K12ac and H4K8/K16ac^30^, suggesting a potential for crosstalk. Thus, understanding how writers and histone modifications influence each other will be very important for understanding how chromatin states are regulated. The histone code hypothesis also supports the idea that histone modifications collectively contribute to transcriptional regulatory logic. However, these are all inherently combinatorial problems that exponentially increase the number of samples and experimental conditions to be tested. Furthermore, it is challenging to determine whether histone residue crosstalk could be achieved by just a single writer protein, as opposed to a situation where a complex of proteins is needed to recognize one modification and then recruit a writer to modify another residue. Here, we investigated whether REMY could map the crosstalk between all possible pairs of acetylated histone residues. In other words, does the absence of acetylation at one site affect the ability of p300 to acetylate another site?

To assess crosstalk between acetylated residues, we measured acetylation at all residues for yeast cells expressing distinct Lys-to-Arg mutants of histones H3 and H4, and p300. Each yeast sample was labeled with anti-C-Myc and residue-specific acetylation antibodies. The samples were gated for cells displaying the histone tail (positive C-Myc signal above unlabeled cells). Then, the relative acetylation level of each residue was calculated using a secondary gate for cells showing labelling with both antibodies (**Figure 4**). First, as expected, the acetylation signal for the mutated residue dropped sharply compared to the wild type peptide. This trend was not seen for H3K4ac (**Figure 4**), but it is important to note that the wild-type histone H3 did not show acetylation at this site.

**Figure 4.**
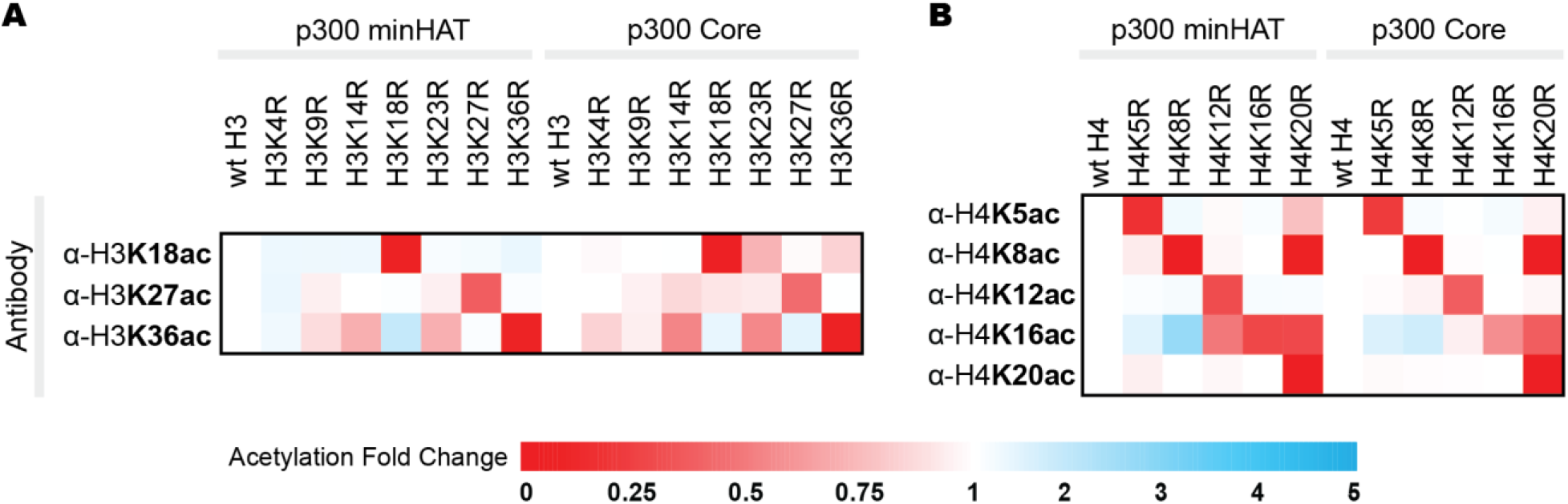
REMY rapidly maps histone residue crosstalk effects on p300 activity. The yeast strain measured is listed across the top of each heatmap and the acetylation site probed by a specific antibody using flow cytometry is listed to the left of each row. Samples were double labeled with anti-C-Myc and anti-acetylation antibodies. An initial gate was created for cells showing C-Myc expression, indicating expression of the histone tail. Within this population, a secondary gate was created for cells also showing fluorescence from the anti-acetylation antibody. The number of cells in the double-labeled gate was divided by the number of cells in the anti-C-Myc gate to quantify the level of acetylation on each histone tail. Acetylation levels were normalized to the acetylation level of the wild-type histone tail for each acetylation site. Each experiment was performed in biological triplicate, having each replicate show the same trend. The average acetylation level from each replicate is shown here. Red is used to indicate a decrease, blue is used to indicate an increase, and white is used to show no change in relative acetylation level at each lysine site.

Interestingly, we observed strong crosstalk between H4K20ac and both H4K8ac and H4K16ac. Mutating H4K20 to an arginine ablated the strong acetylation that was originally observed at H4K8 and H4K16. This crosstalk was slightly stronger for the minHAT domain compared to the Core domain. Conversely, mutating H4K8 or H4K16 to an arginine did not change the acetylation levels of H4K20ac for either the p300 minimal HAT or Core domains. This observation indicates that H4K8ac and H4K16ac written by p300 relies on the presence of H4K20 being acetylated. To our knowledge, this is the first report of this crosstalk interaction. These results show that REMY is a powerful tool for efficient interrogation of crosstalk between histone residues.

## Discussion

We describe a facile, economical, and high throughput platform for chromatin biology termed REMY, which enables post-translational modification and yeast surface display of histone tails. We have shown that REMY can efficiently map the residue specificities of acetyltransferase domains, determine the specificities of commercially available anti-acetylation antibodies, and reveal crosstalk interactions between acetylated histone residues. Importantly, REMY enables analysis of histone modifications without the need for chemical peptide synthesis, chemical modification of enzymes, or production of recombinant proteins, thereby reducing experimental costs and expertise requirements.

Histone residues identified as acetylated by p300 using REMY are consistent with results from prior work, validating our approach (**Table 1**). Importantly, REMY also uncovered new crosstalk between acetylation sites of histone H4. Investigations in an efficient and controlled in vitro setting such as REMY can elucidate the biochemistry of histone modifications, which may be confounded by other factors in an intracellular context. For example, the histone H3 residue specificity of p300 has been reported to be different in COS-7 (African green monkey kidney) cells, HeLa (human epithelioid cervical carcinoma) cells, and IM (male Indian Muntjac skin fibroblast) cells, while the histone H4 acetylation pattern remained the same across cell types tested^28^. By removing other confounding factors, REMY allows for the direct study of the interaction between the epigenome writer and histone substrate^31^. More practically and of immediate relevance, REMY is able to efficiently screen the suitability or specificity of antibodies. This is an important feature as histone antibodies in particular are notoriously promiscuous and improperly validated by many vendors^19,32,33^.

In conclusion, REMY connects the powerful protein engineering and screening abilities of yeast surface display with chromatin biology. Only standard molecular biology, yeast culture, and flow cytometry analyses are required for REMY. Since these are simple, robust, and automatable protocols, REMY provides a potentially democratizing, cost-effective, and time efficient platform for studying chromatin biology.

## Methods

### Creation of histone H3 and histone H4 plasmid families

Three gene fragments containing the N-terminal tail of histone H3 (aa1-45), the N-terminal tail of histone H4 (aa1-26), and the p300 minimal histone acetyltransferase (minHAT) domain along with the appropriate restriction sites were purchased from Genewiz. The histone gene fragments were cloned to an existing pCTCon2 YESS plasmid using XhoI and EcoRI restriction enzymes, allowing them to be transcribed by the GAL1 promoter. The histone tails were individually placed downstream of an endoplasmic reticulum sequence (MQLLRCFSIFSVIASVLA) and upstream of an endoplasmic reticulum retention sequence (FEHDEL). The resulting plasmids containing either the histone H3 tail or histone H4 tail without any acetyltransferase domain were named pAW1 and pAW2, respectively (Supplemental Table 3). In both cases, the histone tail has an N-terminal HA tag and a C-terminal Myc tag to allow for quantification of expression levels.

The p300minHAT domain was cloned into the pCTCon2 YESS plasmid using NcoI and AvrII restriction enzymes, allowing it to be transcribed by the GAL10 promoter. The p300Core domain was amplified via PCR from pRT2, a plasmid previously used in the lab. The p300Core domain was inserted into the pCTCon2 YESS plasmid using CPEC cloning. The resulting plasmids were transformed into electrocompetent NovoBlue *E.coli* and grown on LB plates supplements with carbenicillin. Colonies were selected from these plates, grown in LB ampicillin liquid cultures, purified, and sent to Genewiz for Sanger sequencing. The Sanger sequencing confirmed the successful insertion of the histone tails and p300 domains into the pCTCon2 YESS backbone. The arginine mutations were introduced by purchasing gene fragments from Genewiz and inserting these new genes into the existing pAW1 and pAW2 plasmids using the NheI and AatII restriction sites inserted directly upstream and downstream of the histone substrates.

### Plasmid transformation into EBY100 yeast

EBY100 *Saccharomyces cerevisiae* were transformed with the histone H3, histone H4, and pYD1-mSA^22^ (Addgene plasmid #39865) plasmids using the Zymo Research Frozen-EZ Yeast Transformation II Kit (Catalog# T2001).

### Yeast cell culturing

Yeast were grown in SDCAA media containing dextrose, difco yeast nitrogen base, bacto casamino acids, Na_2_HPO_4_, and NaH_2_PO4•H_2_O in ddH_2_O. Yeast grow in this media for 16-24 hours at 30 °C shaking at 250 rpm. After this growth period, the yeast were induced by a passage into SGCAA at an OD of 1 and kept in this induction media for 16-24 hours at 20 °C shaking at 250 rpm before analysis. SGCAA is similar to SDCAA media except galactose is added at a ratio of 10:1 to dextrose in SGCAA media whereas only dextrose is included in SDCAA media.

### Labeling mSA yeast with biotinylated peptides

EBY100 yeast containing the pYD1-mSA plasmid were cultured in SDCAA media for two passages before induction in SGCAA media supplemented with 1uM D-biotin for 16-24 hours. After induction, 2*10^6^ mSA yeast cells were labeled with 1uM of biotinylated peptide in 100uL 0.1% BSA PBS for 30 minutes, shaking at 500 rpm at 4C.

### Flow cytometry analysis of induced yeast cultures

Freshly induced yeast cultures were processed for flow cytometry by aliquoting 2*10^6^ cells into individual wells of a 96-well plate for antibody labeling. Each acetylation site was labeled in separate samples to ensure that the antibodies did not occlude each other. 50uL of a primary antibody dilution (Supplemental Table 2) was added to each well and allowed to incubate at 4C, shaking at 500-800 rpm for at least 20 minutes. The samples were then washed of unbound primary antibody by resuspension in 200uL of 0.1% BSA 1X PBS, centrifugation at 3000 G for 2 min, and aspiration. The samples were then labeled with a secondary antibody corresponding to the host animal of the primary antibody. The secondary antibodies were added in 50uL aliquots of 1:250 dilutions of their stock concentrations. The secondary antibodies were allowed to incubate with the samples for at least 15 minutes in the dark, at 4C. The secondary antibodies were washed in the same manner as the primary antibodies. The final sample pellets were resuspended in 200uL of 0.1% BSA 1X PBS when the plate was loaded into the flow cytometer (MACSQuant® VYB). Flow cytometry data was analyzed with FlowJo software. All mapping and binary crosstalk experiments were performed in triplicate with three biological replicates showing the same trend. All fluorescent gates were created based on an unlabeled sample of the same yeast strain (with the same bound peptide when applicable), measured on the same day, as the experimental samples.

## Abbreviations

REMY: Rapid interrogation of Epigenome Modifications using the Yeast surface
HAT: Histone acetyltransferase
YESS: yeast endoplasmic reticulum sequestration screening
p300: EA1 binding protein p300

## Author Contributions

AW, BMR and AJK conceived the study. AW planned and performed the wet lab experiments with guidance from BMR and AJK. AW, BMR, and AJK wrote the paper.

## Acknowledgment

This work was supported by the NSF Emerging Frontiers in Research and Innovation program (NSF-1830910) and the NIH (R21EB023377). The pYD1-mSA plasmid was created by the Park Lab at the University of Buffalo and purchased from Addgene (plasmid #39865).

## Availability of Data and Materials

The datasets generated and/or analyzed during the current study are available from the corresponding authors upon reasonable request.

