## Supplemental Info for "REMY: A platform for the rapid interrogation of epigenome modifications on yeast"

**Supporting Information.**

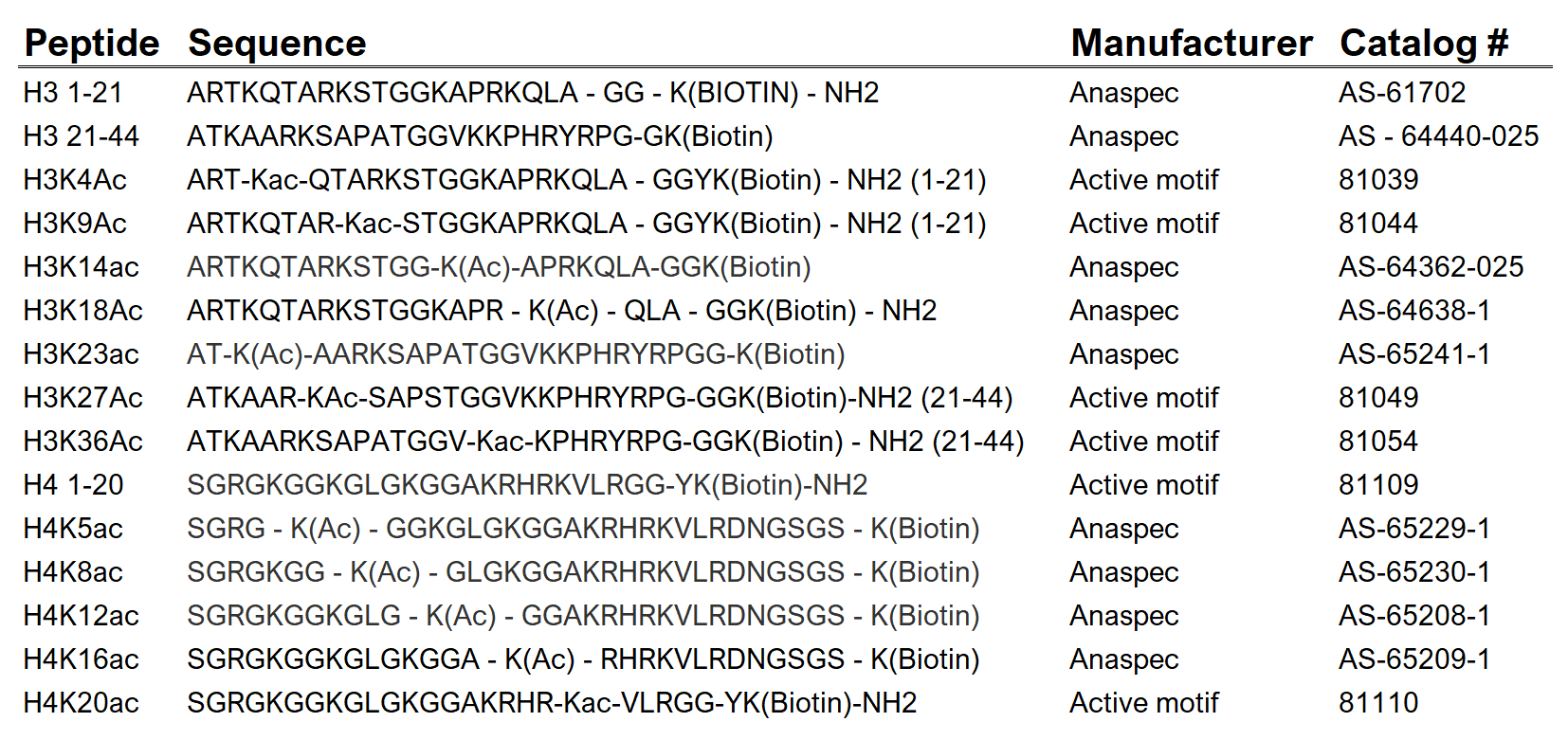
**Supplemental Table 1**. Peptides Used

**Supplemental Table 2**. Primary Antibodies. The antibodies used in Figure 3 to map the acetylation patterns of p300 on the histone H3 and H4 peptide tails are bolded here. The residue specificity tests of these antibodies is shown in the two right most columns. To pass the “synthetic peptide test” the antibody showed binding to the corresponding acetylated histone peptide and no binding to the corresponding unacetylated histone peptide. To pass the “arginine histone test” the antibody showed binding to the wild-type histone peptide when p300 was present and did not show binding to the arginine-mutated histone peptide when p300 was present. For cases where the antibody did not show binding to the wild-type histone in the presence of p300, the arginine histone test was not applicable.

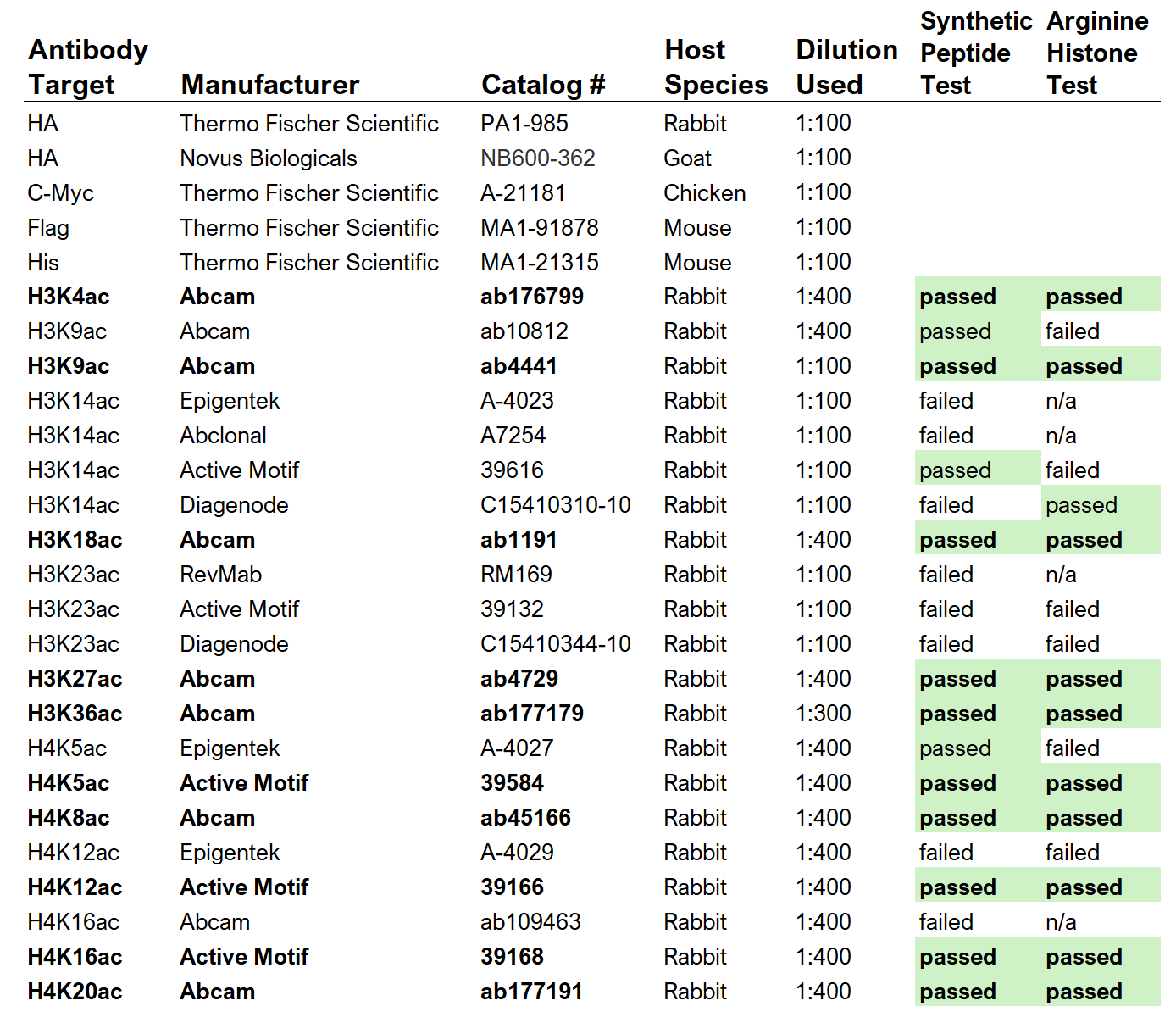

**Supplemental Table 3.** Plasmid Names and Content

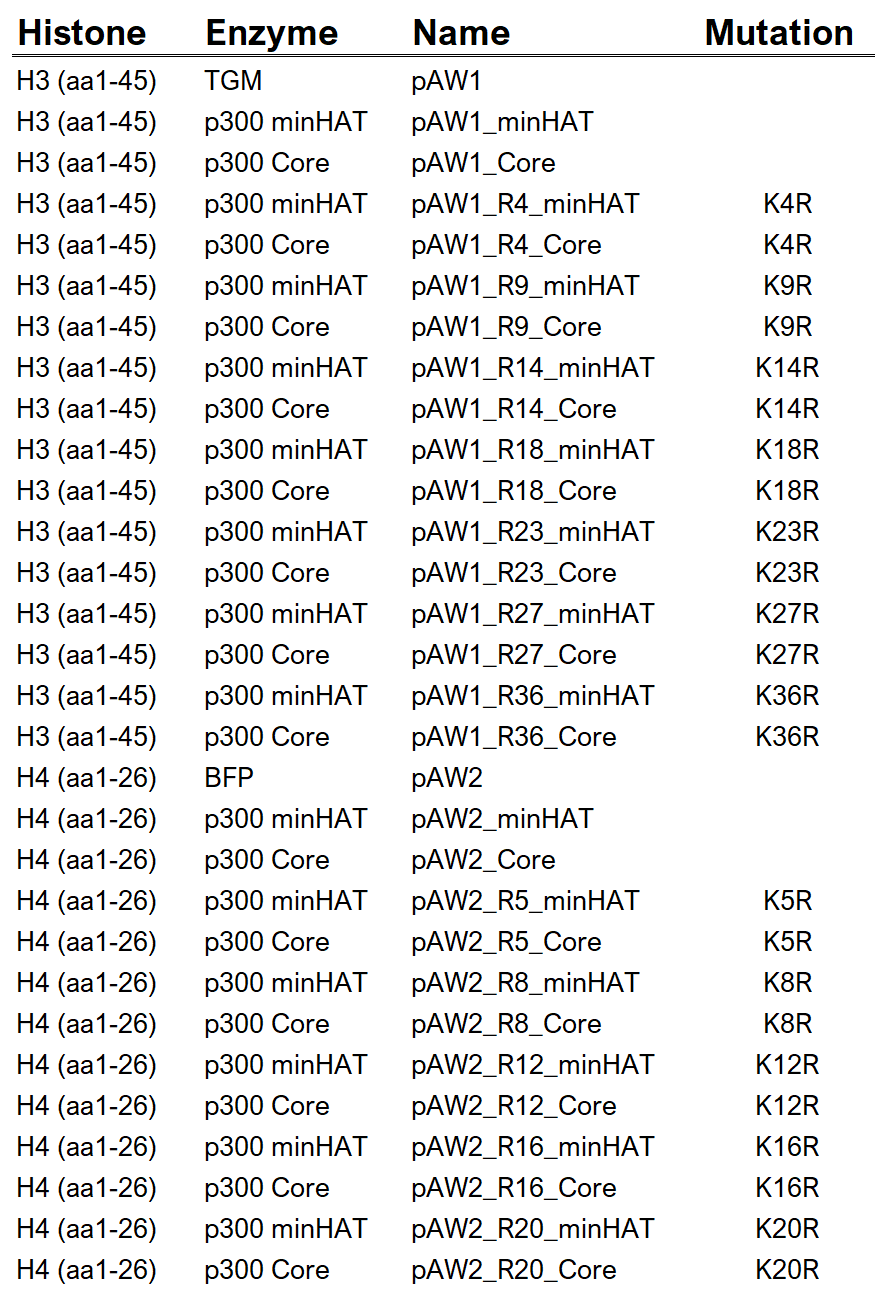

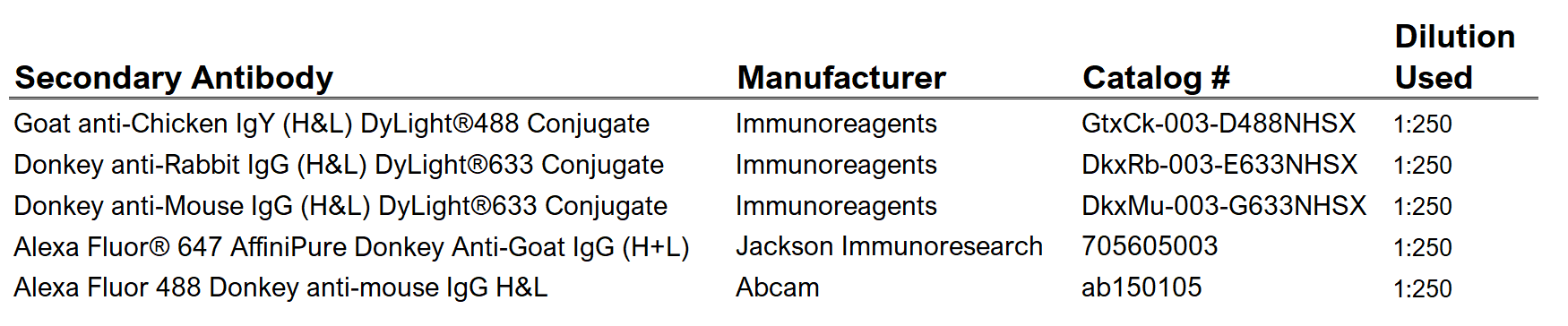
**Supplemental Table 4.** Secondary Antibodies

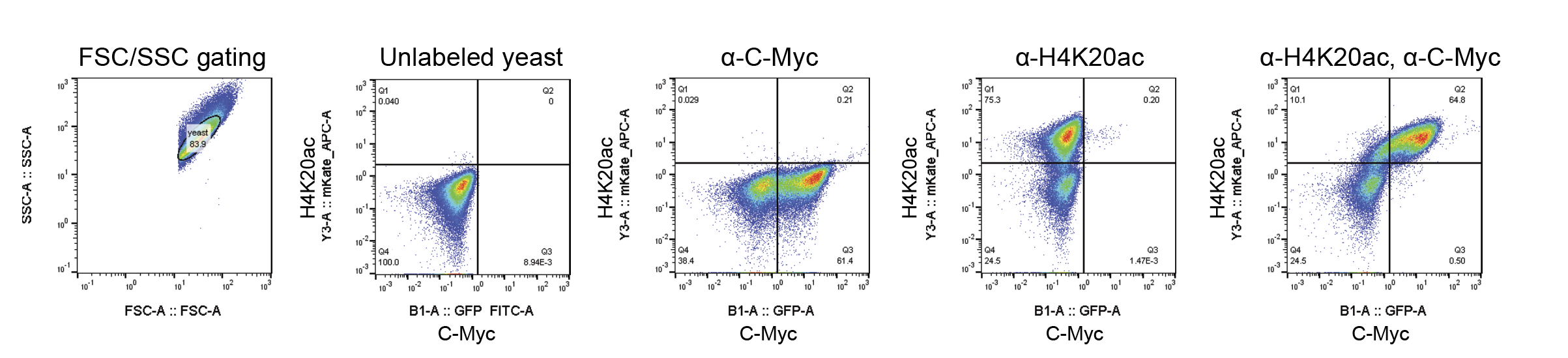

**Supplemental Figure 1**. Flow Cytometry Gating Scheme.

The FSC/SSC gate is based on the expected size of yeast cells. Quadrant gates were created using the unlabeled sample. Acetylation level was calculated based on the number of cells in quandrant 2 divided by the number of cells in quadrants 2 and 3 combined, using the double-labelled sample. All yeast samples shown here are pAW2_R16_minHAT yeast.
